# Beyond translation initiation: dual regulatory interactions with phosphorylated and non-phosphorylated LeishIF4E1

**DOI:** 10.1101/2025.07.16.665100

**Authors:** Durgeshwar Pullaihagari, Nofar Baron, Rajaram Purushostam, Dikla Kamus-Elimeleh, Shiran Dror, Liron Levin, Michal Shapira

**Author notes:** Corresponding Author: Prof. Michal Shapira, Department of Life Sciences, Ben-Gurion University of the Negev, Beer Sheva 84105, Israel.

## Abstract

Translation initiation in eukaryotes begins with the assembly of the cap-binding eIF4F complex at the 5’ ends of mRNAs, through eIF4E. Among the six orthologs of eIF4E in *Leishmania*, LeishIF4E1 is intriguing, as it does not bind any *Leishmania* eIF4G. It is expressed in both life-forms, maintaining efficient cap-binding activity for translation initiation, unlike other LeishIF4Es. We identified two novel phosphorylation sites specifically S108, and S112, along with S203, which is conserved with the mammalian phosphorylation site. A non-phosphorylatable, tagged, mutant of LeishIF4E1 created by substituting the phosphorylated serine residues with alanine [S(108,112,203)A] was generated. The phosphorylation status did not affect the binding of LeishIF4E1 to translation initiation factors. However, the phosphorylated LeishIF4E1 interacted with proteins involved in DNA and chromatin structure, while non-phosphorylated LeishIF4E1 interacted with proteins that assist cells under stress and unfavorable conditions, particularly in gluconeogenesis. This suggests secondary regulatory roles for LeishIF4E1. RNA sequencing of the LeishIF4E1-associated transcripts revealed a tenfold reduction in transcripts binding to non-phosphorylated LeishIF4E1. GO enrichment analysis indicated distinct phosphorylation-dependent transcript specificities for each form. This study highlights the critical role of LeishIF4E1 phosphorylation in modulating the selectivity of protein interactome and transcript associations, extending the role of LeishIF4E1 beyond its well-established function in translation initiation.

**Author Summary:** *Leishmania* are digenetic parasites that transition from sand fly vector to mammalian hosts, undergoing a developmental program of gene expression regulated at the level of translation. Protein synthesis is initiated by the assembly of multi-subunit complexes at the mRNA 5’ cap structure and is recruited onto the ribosomes. There are six cap-binding proteins in *Leishmania* denoted LeishIF4Es1-6, and five LeishIF4G candidates. Among these, the function of LeishIF4E1 is intriguing, as it does not bind any LeishIF4G paralogs, and it is highly expressed in both parasite life forms. We examined how LeishIF4E1 phosphorylation modulates its interaction with proteins and transcripts, given that eIF4E phosphorylation in higher eukaryotes is known to regulate gene expression during malignancy. We identified three phosphorylation sites (S108, S112 and S203) and mutated them to A residues [S(108,112,203)A]. The mutated genes were transfected back into parasites for expression. Using pull-down assays, we identified the interacting proteins and associated transcripts for both the phosphorylated and non-phosphorylated LeishIF4E1 forms. The phosphorylated LeishIF4E1 pulled down proteins involved in DNA and chromatin structure, while the non-phosphorylated LeishIF4E1 interacted with proteins known to assist cells during stress and unfavorable conditions, especially gluconeogenesis. The transcript specificities differed markedly in the number of transcripts bound to both forms, shedding light on the importance of LeishIF4E1 phosphorylation. Furthermore, the phosphorylated form bound more transcripts encoding enzymes related to protein phosphorylation. We also showed that phosphorylation of LeishIF4E1 did not influence the cap-binding ability, as both the forms bind the m^7^GTP cap with parallel efficiencies. Our data suggest that the different cap-binding proteins in *Leishmania* are responsible for different processes and have a transcript-specific effect on gene expression.

## Introduction

Gene expression is regulated at multiple levels, including transcription, RNA processing, and the export of RNA from the nucleus to the cytoplasm for translation on ribosomes. Among these processes, translation enables the most rapid response to changing environments and signals, by selectively recruiting specific mRNAs that produce proteins required for swift and efficient adaptation. Translation of specific mRNAs has also been observed in cells that have undergone malignant transformation [1].

Translation initiation is promoted by assembly of the eIF4F complex at the 5’ cap structure of target mRNAs. This complex comprises the cap-binding protein eIF4E, the scaffold protein eIF4G, which binds to both eIF4E and the RNA helicase subunit eIF4A. In addition, eIF4G interacts with the 40S subunit of the ribosome and the poly(A) binding protein (PABP) at the 3’ end of the mRNA, promoting the circularization of the transcript and increased translation efficiency [2] [3].

Trypanosomatid parasites encode six different cap-binding proteins, denoted LeishIF4Es 1-6, that share limited sequence similarity. LeishIF4Es (1 −6) differ in their cap-binding affinities and perform diverse biological functions [4–9]. In addition, trypanosomatids encode five eIF4G orthologs and three eIF4A homologs [8, 10] [7], generating a variety of eIF4F complexes, each predicted to fulfill distinct cellular roles [11]. Each of the five LeishIF4Gs contains the MIF4G domain, a hallmark of eIF4G proteins [7, 12]. Different LeishIF4Es and LeishIF4Gs assemble into unique complexes with LeishIF4Es (3,4,5 and 6) but not with the LeisheIF4E1 and 2 isoforms [6, 13, 14]. The presence of multiple isoforms of cap-binding proteins and their interacting partners in *Leishmania* and Trypanosomes is particularly intriguing, given that these unicellular organisms do not undergo classical meiosis. Nevertheless, the parasites go through a complex differentiation program, transitioning from promastigotes, which inhabit the intestinal track of sand flies, to amastigotes, as obligatory intracellular parasites, residing within macrophages and cells of the immune system in mammalian hosts [15]. This differentiation is governed by a tightly regulated developmental gene expression program involving specific translation factors.

Recent studies on the *T. cruzi* orthologs of LeishIF4E3 and LeishIF4E4 highlighted that each of these proteins is responsible for translating a specific set of transcripts, promoting parasite survival in the changing environmental conditions [16]. In our recent work, we examined the potential role of LeishIF4E2 and found that this cap-binding protein is involved in cell-cycle progression. Specifically, LeishIF4E2 influences the expression of histone transcripts and the associated stem and loop binding protein, which regulates histone gene expression [13].

Although numerous studies have examined LeishIF4E1 and its *T. brucei* ortholog [17]. The precise function of this protein remains unclear. We have studied LeishIF4E1 extensively, noting that its cap-binding activity persists in axenic amastigotes, a life form that mimics the intracellular form residing within macrophages by exposure to elevated temperatures and reduced pH [6]. Furthermore, we deleted both gene copies of LeishIF4E1 using CRISPR-Cas9 technology and observed that, although the resulting cells are viable, they exhibited changes in their proteome, morphology, and infectivity [18].

Mammalian eIF4E is up-regulated in malignant cells, promoting the formation of lung metastases in a mouse model [19]. However, this effect was suppressed in an S209A mutant of eIF4E, which cannot be phosphorylated at Serine 209 [20]. Studies in eukaryotes have demonstrated that eIF4E is phosphorylated by MNK1/2 at Ser209located, at the C-terminus of the protein [21]. Analysis of knock-in mice in which Ser 209 was mutated to Ala revealed that these mice were resistant to prostate cancer. Furthermore, transcriptome analysis of polysome-associated mRNAs from the eIF4E^S209A^ mutant indicated that eIF4E phosphorylation regulates the translation of specific transcripts [20].

eIF4E is phosphorylated by Mnk1 and Mnk2, both of which are frequently activated in cancer [22]. Accordingly, phosphorylation of eIF4E has been proposed to regulate the expression of proteins involved in cell cycle progression, cell survival, and cell motility. The cap-binding proteins in *Leishmania* are phosphoproteins. LeishIF4E4 is phosphorylated within its extended N-terminal region [22] and LeishIF4E3 is phosphorylated in its non-conserved N-terminus [11] [23] [24]. However, little is known about the phosphorylation of LeishIF4E1 and its potential role during the *Leishmania* life cycle.

In this study, the phosphorylation of LeishIF4E1 was investigated, leading to the identification of three phosphorylation sites, namely, S108,112 and 203. Mutations were introduced to generate a non-phosphorylatable protein (LeishIF4E1^S(108,112,203)A^) and further transfected into WT *L. amazonensis.* This approach enabled examination of how LeishIF4E1 phosphorylation influences the composition of its associated proteome complexes and repertoire of specifically associated transcripts. These findings indicate LeishIF4E1 phosphorylation modulates selective protein complexes and transcript specificity, suggesting an alternate function beyond translation initiation.

## Results

### Structural conservation of the cap-binding pocket and identification of two unique phosphorylation sites in LeishIF4E1 as compared with human IF4E

In higher eukaryotes, phosphorylation of the eukaryotic initiation factor eIF4E occurs as part of the translation regulation process [25]. The structural superposition of LeishIF4E1 (4E1_*L.ama*) and human eIF4E (Human 4E) shows a conservation of their structures, even though the sequence identity is moderate (25.8%). The structural core of the proteins is conserved except for two non-overlapping regions, the N-terminal region, which is not fully visible in the superposition, and a non-conserved region extending from I100 to T126 in LeishIF4E1 (Fig. 1A and 1B).

**Fig 1.**
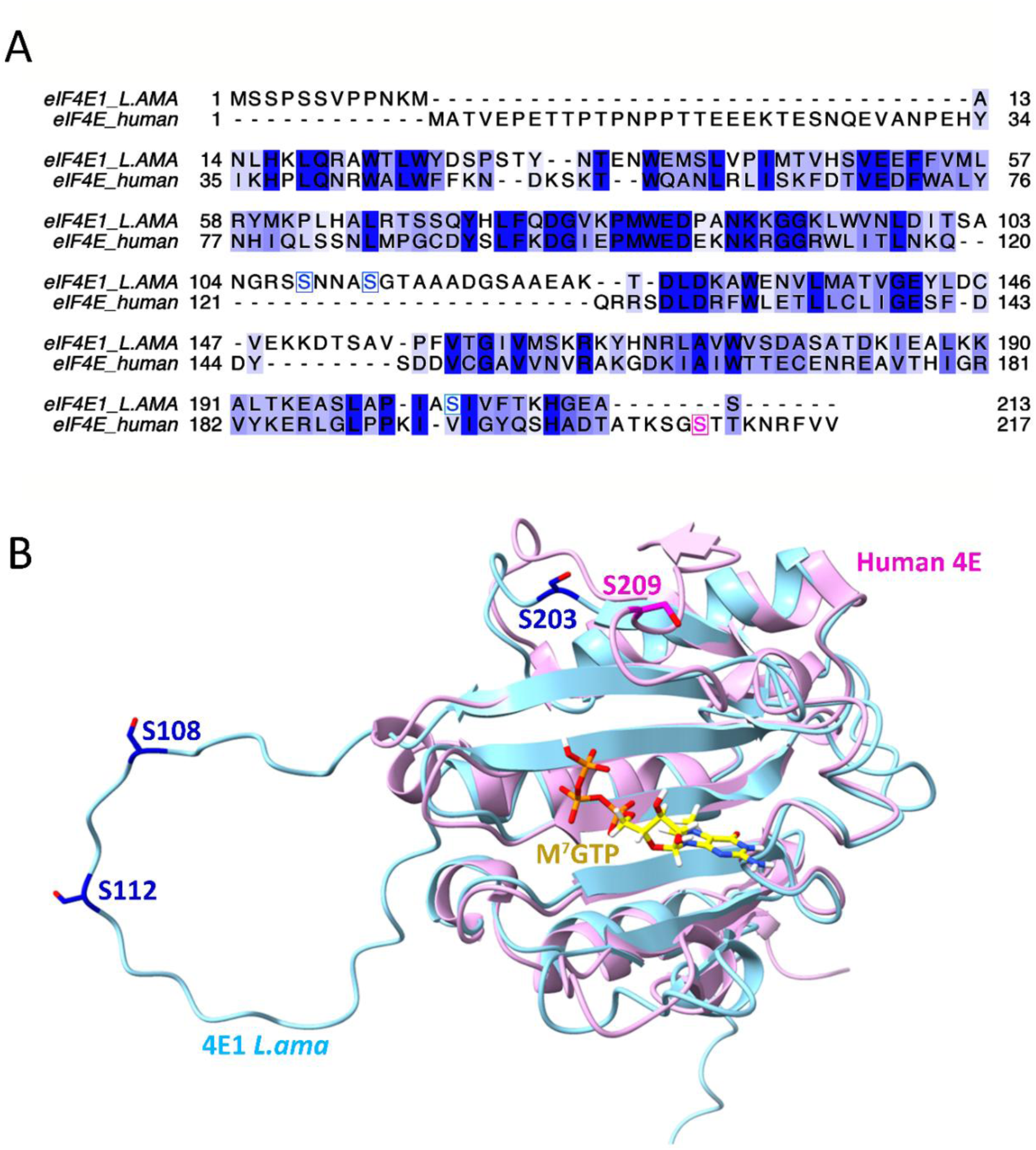
Homology modelling and structural comparison of phosphorylated sites of L. *amazonensis* IF4E1 and human eIF4E. Homology modeling of *L. amazonensis* eIF4E1 (LAMA_000544700) was generated using AlphaFold 3 and compared to the human eIF4E1 structure (PDB code 1ipb) using ChimeraX 1.2.5. **(A)** The amino acid sequence alignment was obtained from Chimera 1.15 and further refined using Jalview (2.10.5). Phosphorylation sites in LeishIF4E1 (S108, S112, S203) are marked in light blue boxes, while the single phosphorylation site in the human eIF4E1 (S209) is marked in magenta. A non-conserved region extending from I100 to T126 in LeishIF4E1 is highlighted by a red box. **(B)** Superposition of LeishIF4E1 (light blue) and human eIF4E1 (light magenta). Phosphorylation sites in LeishIF4E1 are marked in blue, while the single phosphorylation site in the human eIF4E1 is marked in magenta, and the m^7^GTP is highlighted.

Two of the phosphorylation sites, S108 and S112, are located within the non-conserved fragment of LeishIF4E1 extending from I100 to T126. We therefore tested whether phosphorylation in this region could play a regulatory role in modulating LeishIF4E1 function, affecting its interaction with the cap structure and/or with other proteins in its complex (Fig. 1B). We first tested whether the phosphorylation affected its cap-binding activity,and then identified the interacting proteins that were associated with LeishIF4E1 in either of its phosphorylation forms. Finally, we examined how phosphorylation of LeishIF4E1 affected the repertoire of transcripts that bound to LeishIF4E1.

A third phosphorylation site in LeishIF4E1 is located near the C-terminus (S203), in accordance with the phosphorylation site in the human eIF4E1 (S209). In humans, phosphorylated eIF4E had a key role in translation of Myc and ATF4, which promote oncogenic proliferation to regulate the translation of specific transcripts, including Myc and stress-driven oncogenic transcripts in colon rectal cancer cells [20].

### LeishIF4E1 phosphorylation does not impair m^7^GTP binding

To determine whether LeishIF4E1 phosphorylation impacts m^7^GTP binding, affinity assays followed by western blotting were performed. Cell extracts expressing the SBP-tagged phosphorylated or non-phosphorylated forms of LeishIF4E1, were loaded onto m^7^GTP-agarose beads; washed; and eluted by free m^7^GTP. Loaded supernatant (SUP) and elution (ELU) were analyzed via western blotting and quantified by densitometry. As shown in Fig. 2, the E/S ratio revealed no significant difference in m^7^GTP binding between the two LeishIF4E1 forms. These results suggest that the phosphorylated region does not influence cap-binding activity, as it is not located within the m^7^GTP-binding pocket.

**Fig 2.**
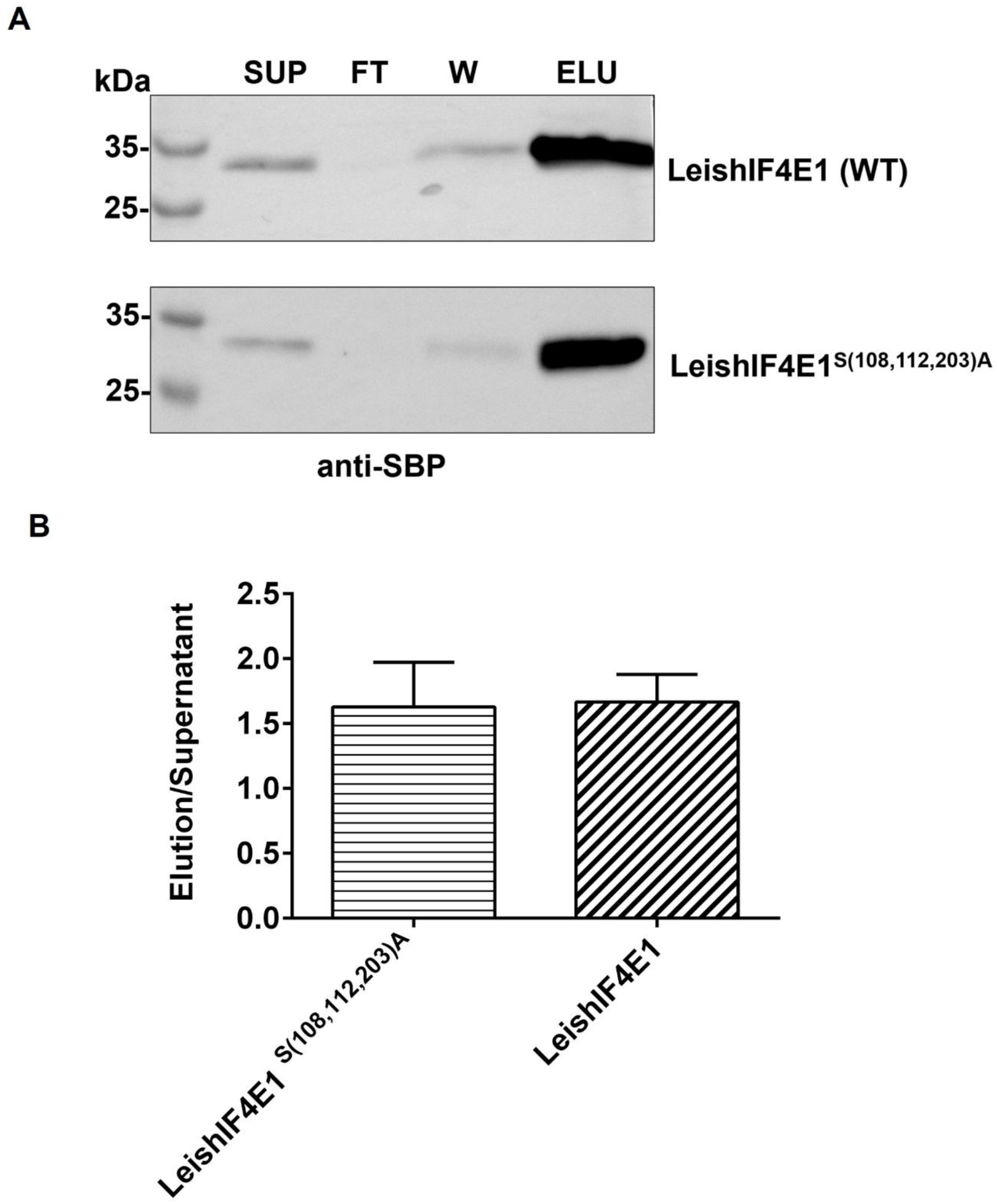
Substitution of the phosphorylated sites in LeishIF4E1 (LeishIF4E1S^(108,112,203)A^) does not affect the m⁷GTP binding activity. Lysates of transgenic cells expressing SBP-tagged LeishIF4E1^S(108,112,203)A^ and the phosphorylated LeishIF4E1 were incubated with m^7^GTP-agarose beads, washed, and eluted with free m^7^GTP. Proteins from the supernatant (SUP, 2%), flow-through (FT, 2%), wash (W, 30%), and elution (ELU, 30%) fractions were separated over 12% SDS-PAGE**. (A)** The gels were blotted and probed with antibodies against the SBP tag. Densitometric analysis was performed on three independent experiments using ImageJ. (B) The ratio of the intensities between elution and supernatant fractions was calculated to evaluate the cap-binding activity.

### LeishIF4E1 phosphorylation status determines distinct protein interactomes

To investigate the role of LeishIF4E1phosphorylation, a mutant cell line that expresses the SBP-tagged non-phosphorylatable LeishIF4E1^S(108,112,203)A^ was tagged with the streptavidin binding peptide (SBP, 4 kD) at its C-terminus and cloned into the pX-transfection vector. The SBP tag facilitated purification of the transgene product using streptavidin-Sepharose beads, enabling us to test how LeishIF4E1 phosphorylation influences the repertoire of its associated proteins and its ability to bind specific transcripts. SBP-tagged luciferase-expressing cells were used as negative control, as previously described [26]. The interactomes of both phosphorylated LeishIF4E1 and non-phosphorylated LeishIF4E^S(108,112,203)A^ were analyzed using mass spectrometry (MS), with significant protein interactions visualized using a Volcano plot (Fig. 3) and detailed in Supp Table 2.

**Fig 3.**
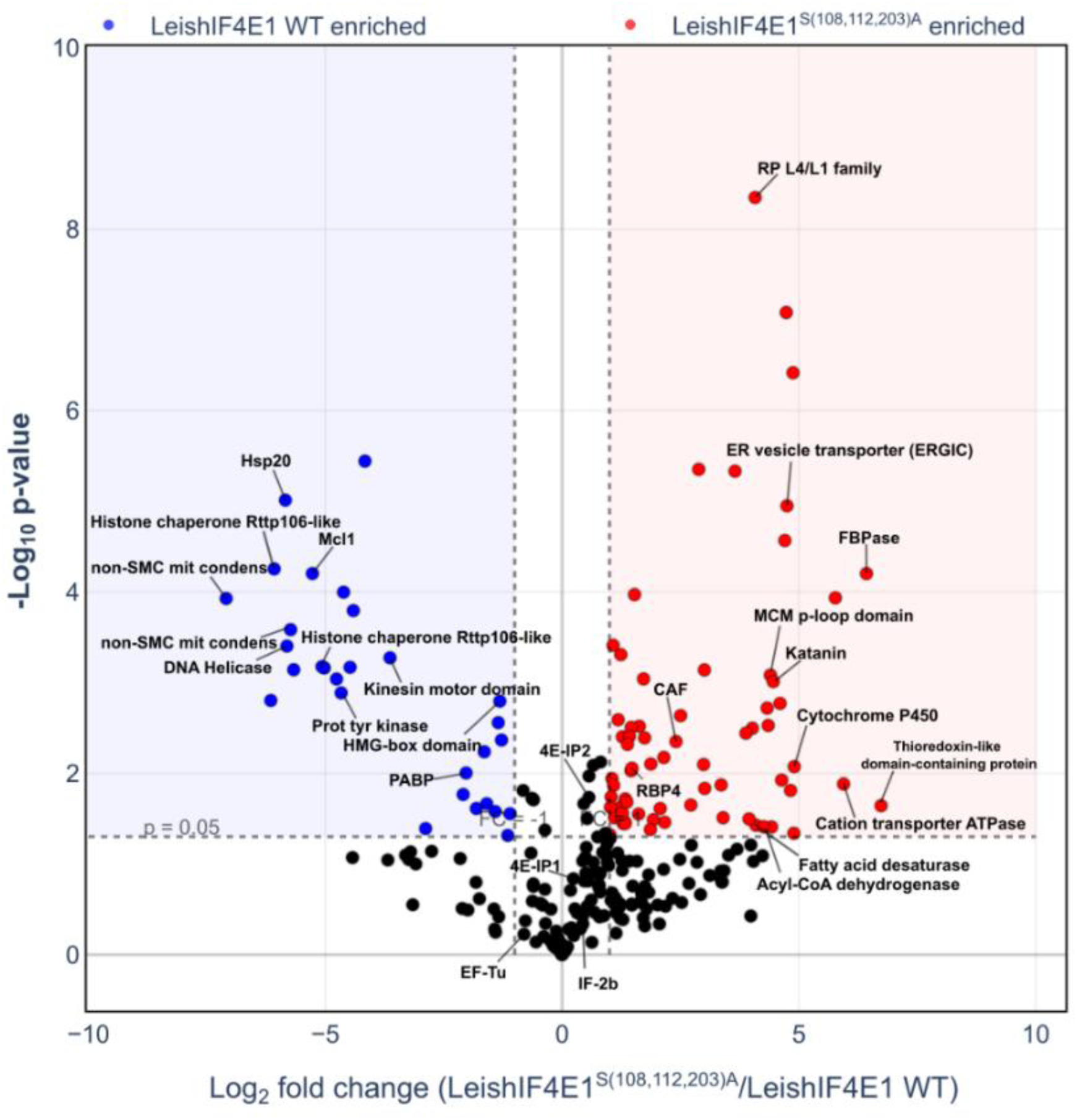
Volcano plots showing proteins that interact with WT-LeishIF4E1 and LeishIF4E1^S(108,112,203)A^. Lysates of cells expressing the SBP-tagged-LeishIF4E1 and– LeishIF4E1^S(108,112,203)A^, along with the SBP-Luciferase control, were pulled down over streptavidin Sepharose and eluted with biotin. The eluents from three independent experiments were subjected to mass spectrometric (MS) analysis. The raw MS data were processed through MaxQuant, and statistical analysis was done through Perseus. The Volcano plot depicts the proteins that were significantly pulled down by the phosphorylated (WT, left) and the non-phosphorylated LeishIF4E1 (LeishIF4E1 S^(108,112,203)A)^ mutant, right). The X-axis represents the log₂ fold change, while the Y-axis shows the log₁₀ p-value for each identified interacting protein. Proteins that were significantly enriched in LeishIF4E1^S(108,112,203)A^ (p-value ≤ 0.05, Log_2_[fold change] > 1) are highlighted in red dots, and those enriched in the WT phosphorylated-LeishIF4E1 are shown in blue dots. Proteins that were not significantly changed are depicted in black dots. The volcano plots were generated using the Plotly library in a custom python script

As shown in Fig. 3, the interactomes of both phosphorylated and non-phosphorylated LeishIF4E1 shared several translation-related proteins, including LeishIF2b (LAMA_000166000), the elongation factor EF-TU (LAMA_000280600), the LeishIF4E-interacting proteins Leish4E-IP1 (LAMA_000806200) and Leish4E-IP2(LAMA_000698700). Leish4E-IP1 specifically interacts with LeishIF4E1 [6] and Leish4E-IP2 that (interacts with several cap-binding proteins [27]. The interactions between the above-mentioned translation-related proteins and LeishIF4E1 indicatedFp that initiation of translation was not affected by changes in the phosphorylation state.

However, the proteins that uniquely interacted with the phosphorylated form of LeishIF4E1 were involved in chromatin organization. Among them were the non-SMC mitotic condensation complex subunit 1 (LAMA_000358900, with an absolute fold change [FC] of 136; LAMA_000129600, FC 53) [28] [29] and the Minichromosome loss protein (Mcl1, LAMA_000192400, FC 38 [30]. Furthermore, enrichment was observed with chromatin organization-related proteins, namely

The interactome of the non-phosphorylated LeishIF4E ^S(108,112,203)A^ mutant showed significant enrichment of proteins involved in metabolic adaptation. A noticeable interactor was Fructose-1,6-Bisphosphatase (FBPase, LAMA_000061200, FC 85), a key enzyme in gluconeogenesis [31] that is typically upregulated under conditions of starvation or nutrient limitation [32]. In addition, proteins involved in fatty acid metabolism and membrane fluidity were detected, such as fatty acid desaturase (LAMA_000010600, FC 28; LAMA_000223500, FC 17) [33] and Cytochrome P450 (LAMA_000181200, FC 29) [34], Lipin, (LAMA_000023900, FC 20) involved in lipid metabolism [35], and Acyl-CoA dehydrogenase (LAMA_000793800, FC 19) [36] [37]. A significant interaction was detected with Thioredoxin-like protein (LAMA_000751400, FC 106), which is crucial for maintaining cellular redox balance [38]. The highly enriched interacting proteins included a group of proteins involved in intercellular transport, the ER vesicle transporter (ERGIC, LAMA_000693900, FC 26) [39];.. [40, 41].

It is important to note that the non-phosphorylated LeishIF4E1 interacted with an alternative subset of proteins associated with chromatin organization, such as the Mini-chromosome maintenance protein 2, MCM, (LAMA_000563800 FC 3.6) [42]; the Histone-binding protein RBBP4 (LAMA_000456000, FC 2.7) [43]; the Chromatin assembly factor 1 (LAMA_000162200, FC 5) [44], and the nucleoskeleton associated protein Katanin_con80 (LAMA_000711200; FC 21) [45]. However, the FC of most of them was lower as compared to the proteins involved in chromatin organization that were bound by the phosphorylated LeishIF4E1.

The phosphorylation state of LeishIF4E1 defines distinct interactomes, highlighting functional divergence. The phospho-mutant primarily associates with proteins involved in metabolic adaptation and oxidative stress management. In contrast, the WT form predominantly interacts with proteins essential for chromatin remodeling, regulation and maintenance of genomic stability. These distinct interactomes highlight the important regulatory role of LeishIF4E1 phosphorylation.

### Phosphorylation of LeishIF4E1 affects the repertoire of associated transcripts

To study the phosphorylation dependent association of LeishIF4E1 with specific transcripts, we repeated the SBP purification of the two LeishIF4E1 forms, phosphorylated and non-phosphorylated, extracted the RNAs that were bound to them and performed an RNA-seq analysis. This revealed major differences in both the number and nature of transcripts associated with each form. A total of 1,845 transcripts were uniquely associated with phosphorylated LeishIF4E1, 184 with the non-phosphorylated form, and 161 with both (Fig 4A).

**Fig 4.**
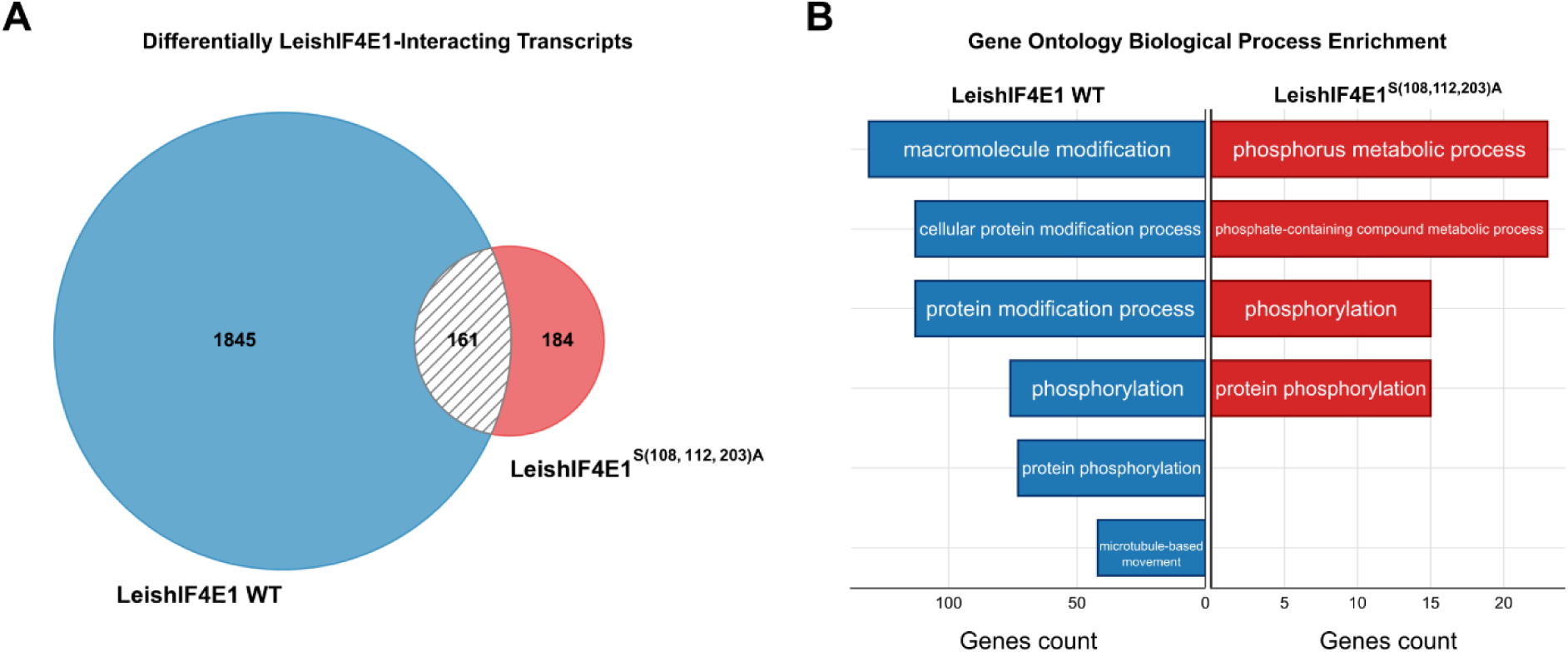
RNA seq analysis of transcripts bound by LeishIF4E1 and LeishIF4E1^S(108,112,203)A^ revealed a fundamental change in the number and identity of bound transcripts. Lysates of cells expressing the SBP-tagged LeishIF4E1 and LeishIF4E1^S(108,112,203)A^ were purified over streptavidin-Sepharose resin, washed and eluted with biotin. mRNAs were isolated from the lysates and from the eluted fractions. Transcripts bound to the LeishIF4E1 WT were compared to LeishIF4E1^S(108,112,203)A^. Enrichment was considered with p-value ≤ 0.05 and log2[fold change]>1. **(A)** A Venn diagram showing the number of associated transcripts, and specific transcripts interacting with LeishIF4E1 WT (1845, blue) or LeishIF4E1^S(108,112,203)A^ (184, red). Shared transcripts (161) are represented as crossed lines. **(B)** Gene ontology biological process (GO BP) enrichment of the associated transcripts was calculated using TriTrypDB (Bonferroni-corrected p-value ≤ 0.05). LeishIF4E1 WT (blue) or LeishIF4E1^S(108,112,203)A^ (red). The X-axis denotes the number of genes associated with each enriched GO BP category. The venn diagram was generated using the Venn library, and histograms were generated by Plotly using available scripts.

A Gene Ontology of Biological Processes (GO BP) analysis was performed (Fig 4B and Suppl Tables 3A (1-6, for analysis of WT cells) and Tables 3B (1-5 for analysis of the mutant cells). In its phosphorylated state, LeishIF4E1 bound several transcripts involved in cell-cycle progression and protein synthesis. These included the cyclin-dependent kinase *crk12,* encoding a protein that is required for transitions through G1/S and G2/M phases, and *rdk1*, encoding a kinase that prevents premature differentiation [46]. In addition, LeishIF4E1 also bound transcripts encoding the kinetochore proteins KKT2 and KKT3, required for chromosome segregation. LeishIF4E1 also bound transcripts encoding cytoskeletal components such as kinesins, dyneins, and FAZ7, which contribute to flagellar movement and attachment. [47, 48].

Although both LeishIF4E1 forms bound transcripts that were categorized as phosphorylation related, their number and identities varied. Among these were 76 transcripts associated with the phosphorylated form of LeishIF4E1, most of them had a general definition as kinases, with no specific targets known. However, three translation-related kinase transcripts were identified, encoding elongation factor-2 kinase-like protein (LmxM.36.4540), *efk-1b,* the isoform-like protein related to elongation factor-2 kinase (LmxM.36.4550) and *eif2ak2* (LmxM.33.2150). In addition, several transcripts were identified as Mitogen Activated Protein kinases (LmxM.13.0440, LmxM.36.3680, LmxM.29.2910, LmxM.31.3250, LmxM.32.2070 and LmxM.36.0720). The Mitogen activated protein kinases comprise a family of protein members, known to convert extracellular stimuli into a wide range of cellular responses and pathways, including cell proliferation, differentiation, motility and survival [49]. The non-phosphorylated form of LeishIF4E1 also bound 3 different Mitogen activated protein kinase transcripts (LmxM.10.0490, LmxM.32.1380 and LmxM.19.0180), supporting its involvement in the variety of cellular processes. However, it should be noticed that the actual binding of specific transcripts by a regulatory protein could have opposing effects, either by enhancing or suppressing their expression. However, the actual binding of these transcripts by LeishIF4E1 indicates that it is involved in key cellular processes.

## Discussion

The role of LeishIF4E1 has remained a subject of considerable interest due to its distinct biochemical properties among the cap-binding proteins encoded in the *Leishmania* genome. Among the six identified cap-binding proteins, only LeishIF4E1 and LeishIF4E2 do not interact with any eIF4G homolog, a crucial component required for translation initiation. While previous studies implicated LeishIF4E2 in regulating the cell cycle progression [13], the precise function of LeishIF4E1 has been less clear. Intriguingly, LeishIF4E1 uniquely maintains active cap-binding ability in axenic amastigotes, differentiating it from its other *Leishmania* orthologs [6]. These observations highlight the unique regulatory potential of LeishIF4E1 and justify further exploration into its function.

Previous studies have indicated that increased expression of eIF4E in eukaryotes enhanced their neoplastic transformation [50]. Further studies documented the role of eIF4E phosphorylation in oncogenic processes [51] [19]. In humans, phosphorylation of eIF4E at S209 by Mnk kinases significantly contributes to tumor progression [50, 52–54]. Knock-in mouse models expressing a non-phosphorylated mutant exhibited resistance to tumor formation, emphasizing the physiological importance of eIF4E phosphorylation in selectively regulating translation of oncogenic mRNAs [21, 55]. These findings highlight the regulatory role of phosphorylation in eukaryotes and support the exploration of similar phosphorylation-dependent mechanisms in *Leishmania*.

We identified three phosphorylated residues in LeishIF4E1, S108, S112, and S203. Notably, residue S203 is located at the C-terminus of LeishIF4E1, analogous to the well-studied phosphorylation site S209 in human eIF4E [51]. However, residues S108 and S112 reside in a region unique to LeishIF4E1, suggesting potential involvement in specific regulatory processes exclusive to *Leishmania*. Nevertheless, the kinase responsible for LeishIF4E1 phosphorylation, the corresponding phosphatase, and upstream signaling mechanisms remain unknown and require further investigation.

Here we showed that phosphorylation of LeishIF4E1 affected its ability to interact with specific proteins and furthermore, with specific transcripts. The statistical analysis of the LeishIF4E1 protein interactome highlighted significant phosphorylation-dependent differences. Notably, the phosphorylated LeishIF4E1 uniquely interacted with chromatin-associated proteins involved in DNA replication, chromatin remodeling, and genomic stability. Most important among these interactors were the histone chaperone Rtt106, essential for nucleosome assembly and heterochromatin silencing, as well the high-mobility group (HMG)-box domain proteins, the non-SMC mitotic condensin complex subunits, and various DNA repair factors. Conversely, the non-phosphorylated form is associated mostly with metabolic enzymes linked to gluconeogenesis and fatty acid metabolism, including Fructose-1,6-Bisphosphatase and Cytochrome P450. These unique interactions indicate phosphorylation-dependent functional specialization of LeishIF4E1 ranging from genomic stability to maintenance or metabolic adaptation.

We expanded the analysis of phosphorylated and non-phosphorylated LeishIF4E1 to identify specific transcripts that bind to each of the LeishIF4E1 forms. Phosphorylated LeishIF4E1 preferentially co-purified with transcripts encoding cell-cycle regulators, including the cyclin-dependent kinase CRK12 [56], the kinetochore proteins (KKT2 and KKT3) [57] and cytoskeletal proteins involved in chromosome segregation and flagellar assembly. In addition, the phosphorylated LeishIF4E1 bound to the transcript encoding RDK1, which is known to be a repressor of differentiation [58] In contrast, the non-phosphorylated form preferentially interacted with transcripts encoding developmental and metabolic regulators, including CRK9, the differentiation-associated kinase RDK2, MAP kinases MPK3 and MPK9 [58], as well as transcripts encoding enzymes involved in nucleotide salvage and inositol signaling pathways. Overall, these findings suggest that phosphorylation of LeishIF4E1 affected the repertoire of transcripts it could bind, supporting its involvement in regulatory activities during distinct cellular states. Furthermore, these transcriptomic results show a correlation with the proteomic data, which show that phosphorylated LeishIF4E1 preferentially interacted with chromatin-associated proteins and factors essential for maintaining genomic stability. The enrichment of chromatin regulators upon phosphorylation likely reflects the cellular requirements for maintaining chromatin structure during DNA replication and active cell proliferation.

It is important to note that despite the changes observed in generating proteomic complexes that were affected by LeishIF4E1 phosphorylation, its ability to interact with several translation initiation factors was irrespective of its phosphorylation status, as observed in Fig. 3. Among them were the elongation factor EF-Tu, the initiation factors IF-2b and IF-2, as well as the ribosomal proteins RPL21 and RPL10. LeishIF4E1 also bound repressors of translation such as Leish4E-IP1 and Leish4E-IP2. In addition to these interacting proteins, we observed that both forms of LeishIF4E1 also bound transcripts involved in the translation process, including the mRNAs encoding the Elongation factor 1-alpha (LmxM.17.0081), RPL2 (LmxM.08.0280), eIF3a kinase and ubiquitin-related enzymes (Suppl. Tables 3A,B). Our results indicate that LeishIF4E1 could have multiple functions in driving translation of specific transcripts, as well as in being involved in cell cycle events.

Our results did not show any effect to LeishIF4E1 phosphorylation on binding to m^7^GTP. This is not surprising, since the phosphorylation sites were identified away from the cap-binding pocket. We could expect that these modifications reduced the flexibility of LeishIF4E1 in its cap-binding pocket, yet this was not observed.

There are contradictory studies that investigated how eIF4E phosphorylation affects its cap-binding activity. In 2010 Sonenberg and colleagues showed that eIF4E phosphorylation promotes tumorigenesis and is associated with prostate cancer progression [19]. The level of eIF4E phosphorylation was further shown to be upregulated in several human cancers [22] [59], promoting transformation, proliferation, apoptosis, metastasis and angiogenesis. In contrast, Scheper and Proud showed reduced affinity of eIF4E for the cap-structure upon phosphorylation at residue 209. They also showed that phosphorylation of eIF4E did not affect its ability to bind 4E-BP1. Since this protein competes with eIF4G for the same binding site in the dorsal part of eIF4E, they assumed that eIF4E did not affect its binding to eIF4G, yet this was not shown directly [60].

Unlike our results for LeishIF4E1, in *T. brucei*, phosphorylation of the eIF4E4 ortholog by CRK1 conversely enhanced its cap-binding activity and its interaction with the parasite eIF4G3 and PABP1 [61]. These contrasting effects underline the differential regulatory impacts of phosphorylation across diverse organisms and cap-binding protein types.

Although our findings show functional consistencies between categories of proteins and mRNAs that bind LeishIF4E1, it is difficult to indicate whether these interactions promote or repress the specific function involved. Further genetic analysis could contribute to resolving this issue.

Phosphorylation of EIF4E4 in *L. infantum* occurs primarily within its non-conserved N-terminal extension and relies on interactions with PABP1 rather than on direct cap-binding or eIF4G3 interaction [62]. Additionally, LeishIF4E3 phosphorylation in *L. amazonensis* is not essential for its interaction with LeishIF4G4 under normal conditions, but its mutant-driven dephosphorylated form disrupted its ability to localize in stress granule localization [11]. Thus, phosphorylation events in Trypanosomatids appear to mediate diverse regulatory roles depending on the biological function and structural features of each cap-binding protein.

Gene expression in pathogenic protozoa of the family *Trypanosomatidae* exhibits several unique features, including the presence of multiple eIF4F-like complexes that contribute to protein synthesis. It was previously shown in eukaryotes that the canonical eIF4F complex which is composed primarily of eIF4E and eIF4G subunits, mediates the selection of mRNAs for translation initiation [3, 63]. In *trypanosomatids*, distinct eIF4E paralogs such as eIF4E3 and eIF4E4 have been implicated in translation. Consistent with their discrete functions, eIF4E3 predominantly bound mRNAs encoding proteins involved in anabolic metabolism whereas eIF4E4 mainly associated with transcripts encoding ribosomal proteins [16, 64]. In addition, recent findings reported that LeishIF4E2 interacts with the stem and loop binding protein 2 (SLBP2) and selectively regulates the translation of histone mRNAs [13, 14]. A similar interaction was reported in *T. brucei* [65].

The current findings indicate that phosphorylation of LeishIF4E1 functions as a fundamental molecular switch, modulating the repertoire of proteins and transcripts associated with this factor, which predominantly maintains its activity in both life stages of *Leishmania*. The phosphorylated form preferentially interacts with chromatin remodeling factors and transcripts involved in cell-cycle progression, protein synthesis, and maintenance of genomic stability. Conversely, the non-phosphorylated form primarily associates with metabolic enzymes and developmental regulators implicated in stress response and adaptation to nutrient limitation. This regulatory flexibility provides Leishmania parasites with an adaptive advantage, thereby maintaining a digenetic switch in life forms at varied temperatures and pH. Thus, we conclude that LeishIF4E1 functions beyond its pivotal role in translation initiation. Its phosphorylation emerges as a central mechanism governing modulating translational and metabolic programs essential for parasite survival and developmental flexibility.

## Materials and Methods

### Cell culture

Axenic promastigotes of wild type (WT) *Leishmania amazonensis* (*L. amazonensis*) strain MHOM/BR/LTB0016 were cultured in Medium 199 media (pH 7) supplemented with 10% heat-inactivated fetal calf serum (FCS, Sigma), 5µg/mL Hemin, 0.1 mM adenine, 40 mM HEPES pH 7.4, 4mM L-glutamine (Sartorius), 100U/mL penicillin and 100µg/mL streptomycin (Sigma) at 25°C.

### Structural comparison of LeishIF4E1 and the human eIF4E with highlighted phosphorylation sites

Homology model of *L. amazonensis* eIF4E1 protein (LAMA_000544700) was generated using AlphaFold 3. The predicted structure was aligned and compared to the human eIF4E1 crystal structure (PDB ID: 1IPB) using ChimeraX 1.2.5. Amino acid sequence alignment was initially conducted with Chimera 1.15 and further analyzed using Jalview (v2.10.5). Key phosphorylation sites (S108, S112, and S203) and the corresponding phosphorylated S209 residue in the human eIF4E1 were highlighted.

### Cloning and transfections

To generate the transgenic cells overexpressing LeishIF4E1 with a streptavidin tag, the ORF from genomic DNA of L. *amazonensis* (LAMA_000544700) was amplified using gene-specific primers and cloned into BamHI/XbaI restriction sites of a pX-based transfection cassette, pX-i-target ORF-i-SBP, where i represents the intergenic region derived from HSP83 of *Leishmania* and SBP is the 4 kDa streptavidin binding peptide tag [66], generating a pX-iLeishIF4E1-SBP plasmid. The plasmid was transfected into mid-log phase *Leishmania amazonensis* cells.

### Identification and mutagenesis of phosphorylated sites

Three phosphorylation sites were identified in LeishIF4E1 based on sequence homology with human eIF4E and verified through mass spectrometric analysis of *L. major* proteins enriched for phospho-proteome. Briefly, the proteins were reduced with 3 mM DTT (60°C for 30 min), modified with 10 mM iodoacetamide in 100 mM ammonium bicarbonate, and digested in 10% acetonitrile and 10 mM ammonium bicarbonate with trypsin (Promega) overnight at 37°C. The tryptic digested peptides were eluted with different gradients of acetonitrile with formic acid in water, dried, and re-suspended in 0.1% formic acid. The resulting peptides were resolved by reverse-phase chromatography. 20 % of the peptides were analyzed by LC-MS/MS, and the remaining 80% of the peptides were enriched for phospho-peptides on titanium dioxide (TiO2) beads. The analysis was done by LC-MS/MS on Q Exactive plus (Thermo) and identification was performed by Proteome Discoverer software version 1.4. Semi-quantitation was done by calculating the peak area of each peptide.

The identified phosphorylation sites were S108, S112, and S203. Site-directed mutagenesis of the three phosphorylation sites was performed using a pairwise PCR reaction using the pUC19 plasmid vector harboring the LeishIF4E1 gene (LAMA_000544700). Three pairs of overlapping primers were designed to introduce alanine substitutions at serine 108,112 and 203, each primer pair containing desired mutation (TCT to GCT for Ser to Ala) (primer sequences provided in Suppl Table 1). Each mutation was amplified independently using the Q5 high fidelity polymerase, and pUC19 LeishIF4E1 as template. Equal amounts (100 ng each) of purified products were combined for overlap extension PCR to assemble the full-length mutant gene. The final PCR product was cloned into pUC19 which was further sequenced and verified for the presence of the mutations. The resulting plasmid bearing the mutations was amplified and cloned into the BamHI/XbaI restriction sites of the pX-based transfection cassette, generating pX-i-LeishIF4E1^S(108,112,203)A^ -i-SBP, where ‘i’ represents the intergenic region derived from HSP83 of Leishmania and SBP is the 4kd streptavidin binding peptide tag and transfected using the above described protocol, and selected for antibiotic resistance against G418 (200 μg/mL).

### *In vivo* protein pull-down and mass spectrometry

Proteins from the transgenic lines expressing the episomal SBP-tagged LeishIF4E1 and LeishIF4E1^S(108,112,203)A^ (the non-phosphorylatable mutant version) were pulled down over streptavidin-Sepharose beads (GE Healthcare). Mid log phase cells (∼2×10⁷ cells/mL; 100 mL) were pelleted down at 4000g for 5min at 4°C, washed twice with phosphate-buffered saline (PBS), and once with PRS buffer containing 35 mM HEPES-KOH, pH 7.5, 100 mM KCl, 10 mM MgCl_2_. Cells were lysed in 1ml of 1% Triton X-100 in PRS^+^ buffer (PRS buffer with the addition of 1 mM DTT, 2 mM iodoacetamide, 7.5 mM sodium fluoride, 50 mM β-glycerophosphate, along with 2X protease inhibitor cocktail) and incubated on ice for 10 min. The lysates were centrifuged at 20000g for 20 min at 4°C. The proteins from the clarified lysates were bound to 200µl of streptavidin-sepharose beads (prewashed in PRS^+^ buffer) by incubating for 1 hour at 4°C while rotating at 10 rpm. The flow through (FT) was collected and the beads were washed 4 times with PRS^+^ (2ml each) and once with 1ml that was collected. Beads were eluted in 500 µl of PRS^+^ buffer containing 5 mM biotin. 200µl of the eluent was sent for mass spectrometry analysis at the Smoler Proteomics Center at the Technion, Israel (https://proteomics.net.technion.ac.il/) where they extracted the proteins using their standard protocols, followed by mass spectrometry using LC-MS/MS on Q Exactive plus (Thermo). Control cells expressing luciferase-SBP were pulled down using a similar protocol.

Raw MS data were analyzed by the MaxQuant software, version 1.5.2.8. The data were searched against the annotated *L.amazonensis* proteins from TriTrypDB (https://tritrypdb.org/tritrypdb/app). The MaxQuant settings selected were a minimum of 1 razor/unique peptide for identification with a minimum peptide length of six amino acids and a maximum of two missed cleavages. For protein quantification, summed peptide intensities were used. The log2 of LFQ intensities were compared between the three biological repeats of each. The three biological repeats of each group were compared by the Perseus software [67].

The proteins pulled down in each of the transgenic lines were compared with those pulled down in the SBP-Luciferase expressing cells, maintaining proteins showing a P value<0.05. Our comparisons were performed on the genome of *L. mexicana*, since its genome is far better annotated than that of *L. amazonensis*, and their genomes are rather close.

### m^7^GTP binding by LeishIF4E1

Mid-log transgenic lines (∼2×10⁷ cells/mL; 50 mL) expressing the Leish4E1-SBP and LeishIF4E1^S(108,112,203)A^ mutant were pelleted down at 4000g, 5min, 4°C, washed twice by resuspending in 1XPBS followed and once with column buffer (CB) containing 20 mM HEPES, pH 7.4, 2 mM EDTA, 1 mM DTT and 50 mM NaCl using similar centrifugation steps as above. Cells were lysed in 1% Triton X100 in column buffer amended with 1 mM DTT, 2 mM iodoacetamide, 50 mM β-glycerophosphate, 7.5 mM sodium fluoride and 2X protease inhibitor cocktail and incubated on ice for 10 min followed by centrifuging at 20000g for 20 min at 4°C. The clarified lysates were incubated for 1 h with 150 μl m^7^GTP-agarose beads pre-loaded on a column and prewashed twice with CB, allowing equilibration. Following binding, the flowthrough was collected, and the beads were washed with CB 3 times. The cap-binding complexes were eluted with CB supplemented with 200 μM free m^7^GTP. The proteins were precipitated with trichloroacetic acid (TCA) at a final concentration of 10% and resuspended in sample buffer. The proteins were resolved over 12% SDS-PAGE gels, blotted, and probed with anti-SBP antibody. Band intensities were measured by densitometry analysis using ImageJ software. The binding was evaluated by calculating the ratio of intensities between the elution and supernatant fractions.

### Extraction of LeishIF4E1 associated transcripts

RNA from the cell lysates corresponding to transgenic lines expressing the Leish4E1-SBP and LeishIF4E1^S(108,112,203)A^ were pulled down over streptavidin-sepharose beads following the protocol performed for proteomics and eluted using 5 mM biotin. The eluent was concentrated by precipitation overnight at −20°C using 0.1:10 (vol/vol) 5M sodium acetate and 100% isopropanol, centrifuged at 12000g for 30 min at 4°C. The resultant pellet was dissolved in 100µl of DEPC-treated water, and the RNA was extracted by adding 4 volumes of Trizol following the procedure mentioned by the manufacturer. Simultaneously, the total RNA from the cell lysates used for LeishIF4E1-associated transcripts was isolated using Trizol.

### RNA-seq libraries

RNA integrity was determined on a QIAxcel device (QIAGEN, QIAxcel RNA QC Kit v2.0), and concentration was determined using a QuantiFluor® RNA System (Promega, #E3310) on a Qubit™ Flex Fluorometer (Invitrogen). Poly A enrichment was conducted from 2µg of total RNA using Dynabeads™ mRNA Purification Kit (for mRNA purification from total RNA preps) following manufactuer guidelines.Stranded RNA-seq libraries were constructed using NEBNext® Ultra™ II Directional RNA Library Prep Kit for Illumina® (New England Biolabs, #E7760). The molarity of libraries was determined using QIAxcel (QIAxcel DNA High Sensitivity Kit) and QuantiFluor® dsDNA System (Promega, # E2670) on a Qubit™ Flex Fluorometer (Invitrogen). Libraries were sequenced (150PE) on a NovaSeq X (Illumina).

### Data Preprocessing and Quality Control

Raw RNA-seq reads were processed using the NeatSeq-Flow platform (Sklarz et al., 2017), a flexible workflow system for high-throughput sequencing. Quality control on both raw and trimmed reads was performed using FastQC (Andrews, 2010), and results were aggregated using MultiQC (Ewels et al., 2016). Adapter and quality trimming were carried out with Trim Galore (Krueger, 2015), using a quality threshold of 25 and a minimum read length of 25 bp.

### Transcript Quantification

Transcript-level quantification was performed using RSEM (Li and Dewey, 2011), utilizing Bowtie2 (Langmead and Salzberg, 2012) for read alignment. A forward strand probability of 0.5 was specified. Prior to quantification, the GFF annotation file from *Leishmania mexicana* (TriTrypDB-57) was preprocessed using the CGAT gff2gff tool (Sims et al., 2014). The annotation was extended by 1000 bp upstream and downstream for each feature using the --flank-method extend strategy to capture regulatory regions relevant for expression estimation. The resulting corrected GFF was used to build the RSEM reference together with the reference genome.

### Differential Expression Analysis

Gene-level differential expression analysis was performed using the NeatSeq-Flow platform DESeq2 (Love et al., 2014) module. Expression matrices from RSEM were used as input. The statistical model included a batch as well as genotype and fraction interaction, predefined contrasts tested fraction comparisons within LeishIF4E1^S(108,112,203)A^ (mutant) and LeishIF4E1(WT) separately. Genes with fold change ≥1 and adjusted P-value <.05 were considered as significantly differentially expressed genes. Significant genes were clustered using the ‘eclust’ function from the factorextra R package (10.32614/CRAN.package.factoextra). Enrichment for Gene Ontology biological processes and KEGG pathways was performed using clusterProfiler v4.0 R package [68].

## Author Contributions

Michal Shapira and Durgeshwar Pullaihagari designed the research; Durgeshwar Pullaihagari, Nofar Baron and Michal Shapira analyzed the data; Durgeshwar Pullaihagari and Dikla Kamus Elimeleh performed the experiments; Nofar Baron analyzed the structure of LeishIF4E1 and the phosphorylation sites, Dikla Kamus Elimeleh generated the non-pohosphorylated LeishIF4E1 mutant. Durgeshwar Pullaihagari, Michal Shapira, Shiran Dror and Nofar Baron wrote the paper. Durgeshwar Pullaihagari, Shiran Dror, Nofar Baron, Dikla Kamus-Elimeleh and Rajaram Purushotham reviewed the manuscript. Figures were made by Durgeshwar Pullaihagari, Nofar Baron and Shiran Dror. Liron Levin analyzed the RNAseq data. All authors have read and approved the final manuscript.

## Acknowledgments

We thank the members of the Smoler Center for Proteomic analysis at the Technion, Israel, for helpful discussions prior to protein analysis. We thank the Kreitmann School for advance Graduate Studies for fellowship support of DKEm NB and PR.

